# Neuroblastoma-derived Extracellular Vesicles Disrupt the Integrity of Central Nervous System Barriers via Tight Junction Modulation

**DOI:** 10.64898/2026.09.06.749705

**Authors:** Tal Manko, Ishai Luz, Meshi Zorsky, Valeria Feinstein, Shirel Ben David, Elie Beit-Yannai, Gad Vatine, Tomer Cooks

**Author notes:** **Corresponding authors:** Tal Manko, The Shraga Segal Department of Microbiology, Immunology & Genetics, Ben-Gurion University of the Negev, Beer Sheva, Israel, Tomer Cooks, The Shraga Segal Department of Microbiology, Immunology & Genetics, Ben-Gurion University of the Negev, Beer Sheva, Israel.

## Abstract

**Background:** Neuroblastoma (NB) is a pediatric malignancy that predominantly affects young children and can be associated with neurological complications even in the absence of direct central nervous system (CNS) invasion. The CNS is protected by specialized endothelial barriers whose integrity depends on tightly regulated intercellular junctions and extracellular matrix homeostasis. Here, we investigated whether extracellular vesicles (EVs) released by NB cells contribute to CNS endothelial barrier dysfunction and examined the underlying molecular mechanisms.

**Methods:** EVs were isolated from early- and late-differentiated murine NB cells and characterized by nanoparticle tracking analysis and transmission electron microscopy. EV protein cargo was analyzed by mass spectrometry-based proteomics. The effects of NB-derived EVs on CNS endothelial barrier function were evaluated using brain microvascular endothelial cell monolayers by measuring trans-endothelial electrical resistance and paracellular permeability. EV-associated proteolytic activity was assessed using gelatinase activity assays and zymography, and the contribution of matrix metalloproteinases (MMPs) was examined using pharmacological inhibition. Statistical analyses included two-tailed Student’s *t*-tests and two-way analysis of variance with appropriate corrections and multiple-comparison testing.

**Results:** NB-derived EVs impaired CNS endothelial barrier integrity, decreasing electrical resistance and increasing paracellular permeability. In contrast, EVs derived from non-tumor astrocytes and fibroblasts supported endothelial barrier resistance. Proteomic analysis identified proteins involved in cell junction organization and extracellular matrix remodeling within NB-derived EVs, including MMP-2 and MMP-9. Functional assays confirmed that NB-derived EVs possess gelatinase activity. Pharmacological inhibition of MMP activity substantially attenuated EV-induced barrier disruption and preserved endothelial junction integrity, supporting a major role for EV-associated proteolytic activity in the observed phenotype.

**Conclusions:** Our findings identify NB-derived EVs as mediators of CNS endothelial barrier dysfunction and link their barrier-disruptive effects to EV-associated metalloproteinase activity. These results reveal a potential mechanism of tumor–CNS communication and suggest that targeting EV-associated proteolytic activity may represent a strategy for preserving CNS barrier integrity in neuroblastoma.

**Plain English summary:** Neuroblastoma is a childhood cancer that develops from immature nerve cells. Although it rarely spreads directly to the brain, children with neuroblastoma can experience neurological complications. How neuroblastoma cells communicate with and affect the central nervous system is still not fully understood.

Cells release tiny membrane-bound particles that carry biological material and allow them to communicate with other cells. Cancer cells can use these particles to influence tissues far from the original tumor. In this study, we investigated whether particles released by neuroblastoma cells can affect the protective cellular barriers surrounding the central nervous system.

We found that particles released by neuroblastoma cells weakened the connections between the cells that form these barriers, making the barrier less effective. In contrast, particles released by non-cancerous supporting cells did not cause this damaging effect. We also found that neuroblastoma-derived particles carry active protein-degrading enzymes called matrix metalloproteinases. Blocking the activity of these enzymes substantially protected the barrier from the effects of the cancer-derived particles.

Our findings suggest that neuroblastoma cells may influence the central nervous system without directly entering it, by releasing particles that travel between cells and weaken its protective barriers. Understanding this form of communication may help explain some neurological complications associated with neuroblastoma and could eventually identify new ways to protect the central nervous system from cancer-related damage.

## Background

Neuroblastoma (NB) is the most common extracranial solid tumor in pediatric patients and arises from the developing sympathetic nervous system (SNS). NB tumors can arise throughout the sympathetic nervous system, including the cervical, paraspinal, and celiac ganglia and the adrenal glands[1]. Approximately 65% of NB tumors arise in the abdomen, with half of those localized to the medulla of the adrenal gland[2]. However, NB can also occur in the neck, chest, pelvis, and ∼1% of patients have no detectable primary tumor at the time of diagnosis[3]. Due to the high variability of NB manifestation, clinical signs and symptoms can range from a benign palpable mass with distension to major illness, long-term neurological damage, and death due to substantial tumor spread[4]. The general outcome in NB patients has improved steadily in recent decades, with 5-year survival rates rising to 74%[1]. The improvement is attributed to higher cure rates in the low-risk group, with up to 92% overall survival. However, 50-60% of individuals in the high-risk group will experience relapse[1]. In this group, the 5-year survival rate decreases to approximately 20%, despite more advanced therapies[5].

Extracellular vesicles (EVs) are heterogeneous nano-sized particles ranging from 50 to 200 nm that are secreted by all cell types[6]. The molecular cargo of EVs may include lipids, proteins, and various forms of RNA and DNA[7]. In recent years, EVs have become a significant focus of cell-to-cell communication research, particularly in the fields of cancer progression and metastasis[8,9], diagnosis[10] and drug delivery[11]. Despite this, few studies have delved into the potential roles of EVs in NB. Some reports indicate that NB-EVs may play an essential role in tumor microenvironment (TME) modulation[12]. It was shown that EVs from NB affect monocytes by secreting EVs containing miR-21, thus increasing the levels of miR-155 in the reprogrammed monocytes, which, in turn, secrete EVs back to the NB cells, resulting in the stimulation of cellular inflammatory pathways[12,13]. The released miR-155 is shown to target TERF1, which inhibits telomerase activity, thus increasing telomerase’s non-canonical roles of drug resistance[12]. NB-derived EVs were also reported to affect the TME by decreasing NK cell numbers and promoting the activation of tumor-associated macrophages, thereby creating immunosuppressive conditions. Such interactions in the TME were suggested to mediate resistance of the NB tumors to second-line treatments such as dinutuximab[14].

In health, the blood-brain barrier (BBB) forms as a neurovascular unit that shields the central nervous system (CNS) from potential toxins in the circulation by forming functional tight junctions (TJs) between brain microvascular endothelial cells[15,16]. In turn, the BBB limits paracellular cell migration and passage of most molecules. The integrity of the BBB is disrupted in both primary and metastatic brain tumors, making it more permeable, often referred to as the ‘brain-tumor-barrier’ (BTB)[16]. Neurological complications can occur in NB through several mechanisms, including opsoclonus-myoclonus-ataxia syndrome (OMAS), which occurs in 2-3% of all NB cases[17], long-term neurocognitive impairment[18], intraspinal tumor extension[19–21], and, more rarely, CNS metastasis. However, whether tumor-derived factors can directly influence CNS endothelial barrier function remains poorly understood[22–24].

Despite growing recognition of EVs as critical mediators of tumor–host communication, the mechanisms by which NB-derived EVs affect CNS barrier integrity remain poorly defined. To address this knowledge gap, we investigated EVs derived from early- and late-differentiated murine neuroblastoma cell lines and assessed their impact on CNS endothelial barrier function using an *in vitro* CNS endothelial cell model. Here, we demonstrate that NB-derived EVs are enriched in TJ-associated proteins and contain matrix metalloproteinases (MMPs) together with gelatinolytic activity. Functionally, these EVs induce a marked and sustained disruption of endothelial barrier integrity, an effect that is reversible upon pharmacological inhibition of metalloproteinase activity. Together, our findings identify NB-derived EVs as active effectors of CNS barrier dysfunction and uncover a mechanistic link between EV cargo composition and endothelial TJ destabilization.

## Methods and Materials

### Study design

This in vitro study investigated the effects of EVs derived from early- and late-differentiated murine neuroblastoma cells on CNS endothelial barrier integrity. EVs were characterized by nanoparticle tracking analysis (NTA) and transmission electron microscopy (TEM), their protein cargo was analyzed by proteomics, and their effects on endothelial barrier function were assessed using TEER and paracellular permeability measurements. EV-associated proteolytic activity and its contribution to barrier dysfunction were evaluated using gelatinase assays, zymography, and pharmacological MMP inhibition.

### Cells

All cell lines were grown in Dulbecco’s Modified Eagle Medium (DMEM) with 10% fetal bovine serum (FBS), 1% L-Glutamine (L-Glu), and 1% penicillin G and streptomycin (PENSTREP). Cells were grown in 6-, 10-, and 15-cm Petri dishes; when the cells reached 80-90% confluency, the media was removed from the dishes, and phosphate-buffered saline (PBS) was introduced against the dish walls to clean the residual media. PBS was removed from the dish, and trypsin was added to the dish for 5 minutes of incubation at 37 °C in a humid incubator with 5% CO_2_. After the set time, DMEM was added twice the amount of trypsin, and the cells were transferred to 15 mL tubes and centrifuged at 1,500 RPM for 3 minutes. Pellets were resuspended in DMEM, and a portion of the cells were transferred to a new DMEM-containing culture dish.

### EV isolation

A standard serial ultra-centrifugation method was used to isolate EVs. Briefly, cells were grown in a 150 mm cell culture plate (Biofil, Guangzhou, China) to reach 80-90% confluency. 100 mL of medium was collected from 4 Petri dishes and centrifuged at 1,500 RPM for 3 minutes. The supernatant was collected and centrifuged at 15,000 × g for 30 minutes. After removing the cell debris, the media was passed through 0.22 µm filters. The media was then centrifuged at 100,000 × g (38,700 RPM) for 90 min at 4 °C. The pellet obtained was resuspended in 5 mL of 1X PBS, and all relevant samples were recollected for another round of 100,000 × g for 90 min at 4 °C. The supernatant was discarded, and the pellet was resuspended in 1X PBS.

### EV size exclusion

For size-exclusion chromatography (SEC), EVs isolated by ultracentrifugation were passed through a 70 nm column (qEV, Izon Science) according to the manufacturer’s guidelines. EVs were collected from fractions 4-6 and taken for TEM for further analysis.

### Nanoparticle tracking analysis (NTA)

NTA measurements were performed using a NanoSight NS300 Instrument (NanoSight NTA 2.3) (Salisbury, UK) following the manufacturer’s instructions. Samples were processed in duplicates and diluted with PBS (1 mL of EVs pellet was further diluted at a 1:10 ratio) before analysis. NTA post-acquisition settings were optimized and kept constant between samples. Five 60-second-long videos were recorded per sample and analyzed to obtain the mean and mode particle sizes, along with an estimate of the number of particles/mL. A formula was created to calculate the total number of particles obtained from the EVs isolation.

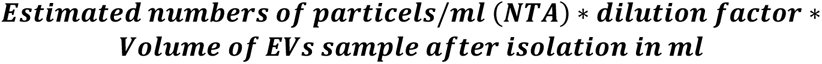

### Transmission electron microscopy (TEM)

EVs were isolated from the NB cancer cell line by utilizing ultracentrifugation followed by SEC. FEI Tecnai T12 G2 TWIN transmission electron microscope (Hillsboro, OR, USA) operating at 120 kV was used. The images were taken with a Gatan 794 MultiScan CCD camera. Briefly, the samples were prepared as follows: 2.5 µL of the sample was applied to a 300-mesh copper grid, and the excess liquid was blotted with filter paper after 1 min. The grid was dried in air for 1 min, then 5 µL of 2% uranyl acetate was applied for negative staining to increase sample contrast. Next, the grid was blotted once more to remove the excess uranyl acetate. Finally, the grid was dried in the air before insertion into the microscope.

### Micro BCA protein measurements

To determine EV protein concentration, EV samples were analyzed using the Micro BCA Protein Assay Kit (23235, Thermo Scientific, USA). EVs were lysed in a mild lysis buffer containing 50 mM Tris-HCl, 150 mM NaCl, 5 mM CaCl₂, and 0.1% Triton X-100 in ultrapure water. We performed the Micro BCA assay according to the manufacturer’s protocol and incubated the plate for 2 h. We measured absorbance at 562 nm using a Multiskan SkyHigh Microplate Spectrophotometer (A51119700C, Thermo Scientific, USA). Protein concentrations were calculated based on a BSA standard curve.

### Zymography

For gelatin zymography, EV samples containing 15 µg total protein were mixed with 4× non-reducing sample buffer without β-mercaptoethanol and not boiled before electrophoresis. Samples were separated on 10% SDS-PAGE gels polymerized with 0.1% gelatin at 110 V for 2.5 h at 4 °C. Following electrophoresis, gels were washed twice in 2.5% Triton X-100 for 15 min per wash to remove SDS and allow protein renaturation. Gels were then incubated in developing buffer containing 50 mM Tris, 200 mM NaCl, 5 mM CaCl₂, and 0.02% Brij-35 (pH 7.6) for 30 min at 37 °C with agitation. The buffer was subsequently replaced with fresh developing buffer, and gels were incubated for an additional 16 h at 37 °C. Gels were stained with 0.5% (w/v) Coomassie Brilliant Blue R-250 in 40% methanol and 10% acetic acid for 30–60 min at room temperature with agitation, then destained in 40% methanol and 10% acetic acid until clear gelatinolytic bands became visible against the blue-stained background.

### ImageJ

Zymography images were converted to 8-bit and inverted using ImageJ to facilitate visualization and densitometric analysis of gelatinolytic bands.

### EVOM2 for TEER measurements of bEND3

bEND3 cells were seeded on 12-well inserts (Greiner, 665640), with 1 mL of DMEM in the insert and 2 mL of DMEM in the well, with 10,000 cells/cm^2^ in each insert. Four hours before seeding, add 150 µL of collagen IV + fibronectin (C5533 and F1141, 4:1 ratio, respectively; Sigma-Aldrich, St. Louis, MO, USA) to the insert to support bEND3 TEER integrity, then remove it before seeding bEND3 cells. Using the patch clam of Epithelial Volt/Ohms for the TEER meter, insert the STX2 electrode in a 90° angle inside the insert with the short electrode and the longer electrode in the well, making sure the electrode doesn’t touch the bottom of the well and insert, waiting until the TEER present on the EVOM2 screen stabilizes, each well - insert being measured twice in each of the three sections of the inserts. Each group was assessed in triplicates, yielding 18 measurements per day per group; after two days following the TEER peak levels, the media were changed, and NB cells and EVs were introduced. bEND3 cells were co-cultured with NB cells in a 1:1 ratio, while bEND3 cells were treated with 100 EVs/cell daily. We introduced EVs from a fresh batch on days 2 and 5. TEER values were presented as Ω·cm^2^ following the subtraction of an empty transwell and multiplication by 1.13 cm^2^ to account for the surface area.

### Proteomics

Four independent biological replicates of N2a and NS20-derived EVs were subjected to mass spectrometry-based proteomic analysis. The EV proteins were cleaved with trypsin and analyzed by the LC/MSMS using the Exploris 480 (Thermo Scientific, USA) mass spectrometer. We analyzed the data using MaxQuant 2.1.1.0 for identification and quantification against the mouse UniProt database with a 1% FDR threshold. We performed proteomic analysis using STRING and Cytoscape version 3.10.1. We generated Venn diagrams using Venny 2.1.0.

### MMPs activity assay

To evaluate MMP activity in NB EVs, we used the EnzChek™ Gelatinase/Collagenase Assay Kit (250-2,000 assays, E12055, Thermo Scientific, USA). Using 96-well glass-bottom black plate (P96-1.5H-N, Cellvis) for fluorescence read, each well contained 200 μL of the following: Negative control 10 µL of DQ and 190 µL of reaction buffer, Positive control 10 µL of DQ, 90 µL of reaction buffer, 100 µL of clostridium collagenase (0.02 U/mL final concentration in well), EVs samples contained 10 µL DQ, 84/88 µL of EVs samples (N2a and NS20 respectively, EVs were suspended in reaction buffer at the final stage of EV isolation) and 106/102 µL of reaction buffer (N2a and NS20 respectively). Plate readings were measured using a Tecan Infinite F200 Pro microplate reader with Tecan i-control version 2.0.10.0 software; temperature range: 36.5 - 37.5 °C; preincubation for 5 minutes comprised 3 minutes of orbital shaking at 3 mm and 2 minutes of waiting. Reading was measured at 495 excitation and 515 emission wavelengths, with a manual gain of 80, 15 flashes, and a single read for each well.

### Statistical analysis

Data are presented as the mean ± standard error of the mean (SEM) from three independent biological replicates (N=3) per experiment, unless otherwise stated. Statistical analyses were performed using GraphPad Prism software. Two-tailed Student’s *t*-tests were used for comparisons between two groups. For comparisons across multiple groups and time points, we performed a two-way repeated-measures ANOVA, with Time as the within-subject factor and Group as the between-subject factor. Matched measurements were stacked into sub columns to account for repeated observations per well. Because sphericity could not be assumed, we applied the Geisser–Greenhouse correction. We performed post-hoc comparisons using Tukey’s multiple-comparisons test, evaluating group differences at each time point (row-wise comparisons, one family per row). Statistical significance was indicated as follows: *p < 0.05, **p < 0.01, ***p < 0.001, ****p < 0.0001. All TEER experiments used 3 biological replicates and 6 reads per biological replicate per group to ensure robustness and reproducibility.

## Results

### Characterization of EVs derived from early and late differentiated NB cells

To study EVs in the context of NB, we used two murine cell lines derived from the mouse neuroblastoma model C1300.[25–27]. The N2a cells represent early-differentiated neuroblast cells [28], while the NS20 cells are more mature neuroblast derivatives[29]. EV characterization was performed following the guidelines recommended in the Minimal Information for Studies of Extracellular Vesicles (MISEV 2023)[30]. Thus, we used nanoparticle tracking analysis (NTA) to determine EV concentration and size (Figure 1A), as well as transmission electron microscopy (TEM) to assess morphology (Figure 1B) [30]. N2a EV concentration resulted in 1.192·10^10^ ± 2.29·10^8^ particles/mL while NS20 EVs concentration was 1.13·10^10^ ± 0.995·10^8^ particles/mL (Figure 1C). The mode size of EVs was 143.87 ± 2.64 nm for N2a EVs, while NS20 EVs were significantly larger at 155.67 ± 3.39 nm (Figure 1D). Using the number of cells on the day of EV isolation, we calculated the particle-to-cell ratio after 5 days of culturing for each cell line, resulting in N2a producing 76.47 ± 1.48 EVs/cell, while NS20 produced significantly fewer at 55.57 ± 0.48 EVs/cell (Figure 1E). Our results show no significant difference in total EV concentration, whereas mode size was significantly larger for NS20 EVs. EV/cell ratios were substantially in favor of N2a EVs, indicating that N2a cells produce more, but smaller, EVs per cell compared to NS20 cells.

**Figure 1.**
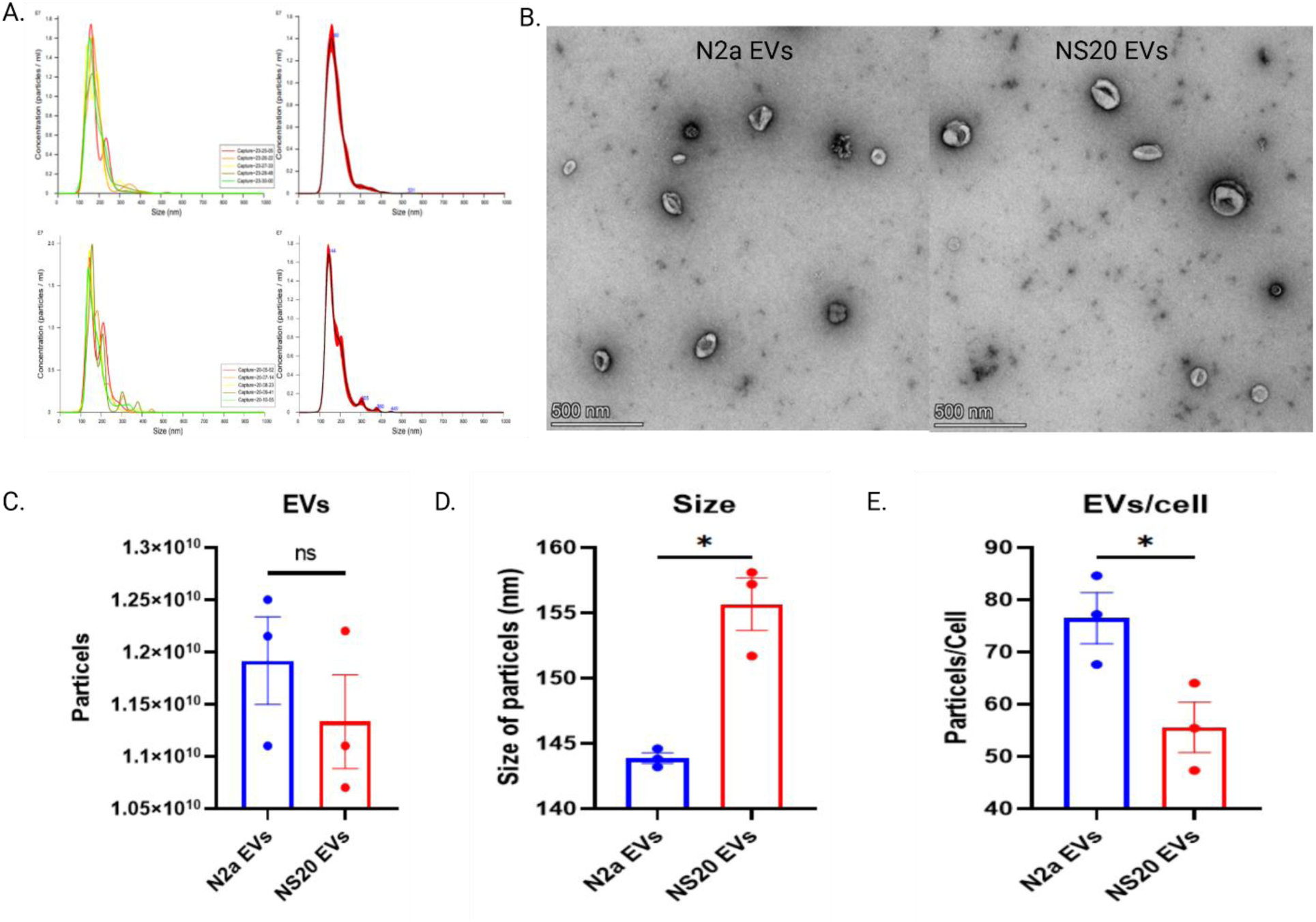
Characterization of EVs derived from early- and late-differentiated NB cells - NTA analysis via NanoSight. NS300 of N2a (top) and NS20 (bottom) (A). TEM imaging of N2a and NS20 EVs (B). Number of EVs from N2a and NS20 cell lines (C). Mode size of N2a and NS20 EVs (D). EVs per cell of N2a and NS20 (E). Biological replicates (n = 3 per group) were used for statistical comparison, with technical replicates averaged to generate a single value per biological replicate. Differences between N2a-derived EVs and NS20-derived EVs were assessed using an unpaired two-tailed Student’s t-test, with p < 0.05 considered significant. Data are presented as mean ± SEM.

### NB-EV proteomes are enriched in proteins involved in cell junction organization and adhesion

To further delineate the composition of the vesicular cargo, NB-EVs from both NB cell lines were subjected to proteomics analysis, yielding 3883 proteins from NS20 EVs and 4271 from N2a EVs. Of the total proteins, 3734 proteins were shared between NS20 and N2a, comprising 96% of the total protein found in NS20 EVs (Figure 2A). EV shared proteomes from NS20 and N2a cells exhibit consistent protein abundance patterns across four biological replicates, shown as log_2_ values (n = 4) (Figure 2B). The average log_2_ data for the shared proteins were calculated and plotted in a heatmap (Figure 2C). A cut-off of log_2_FC > 1 was applied, resulting in 794 significantly elevated proteins in NS20 EVs compared with 508 in N2a EVs (Figure 2D). Proteomics analysis using STRING with a confidence score of 0.7 and the Signal Score biological process visualizer showed that the significantly enriched proteins in N2a EVs were involved in RNA processing (Supplementary Figure 1A), whereas those in NS20 EVs were involved in EV biogenesis and transport (Supplementary Figure 1B). In addition, the signal score molecular function visualizer showed that N2a EV-enriched proteins are mainly involved in RNA binding (Supplementary Figure 1C), whereas NS20-enriched proteins are involved in cell adhesion, extracellular matrix remodeling, cytoskeletal organization, and intracellular transport (Supplementary Figure 1D). Lastly, the signal cellular component visualizer shows that N2a EV-enriched proteins are associated with the nucleosome and the chromosome (Supplementary Figure 1E), while NS20 EV-enriched proteins are associated with the ESCRT system, cell junctions, and the extracellular matrix (Supplementary Figure 1F). These findings coincide with previous work on NB EVs, in which NB proteomics of EVs from human cell lines, primary cells, and bone marrow metastasis patient-derived cell lines has identified proteins associated with neuronal development, cell adhesion, cell junctions, and protein binding [31–33].

**Figure 2.**
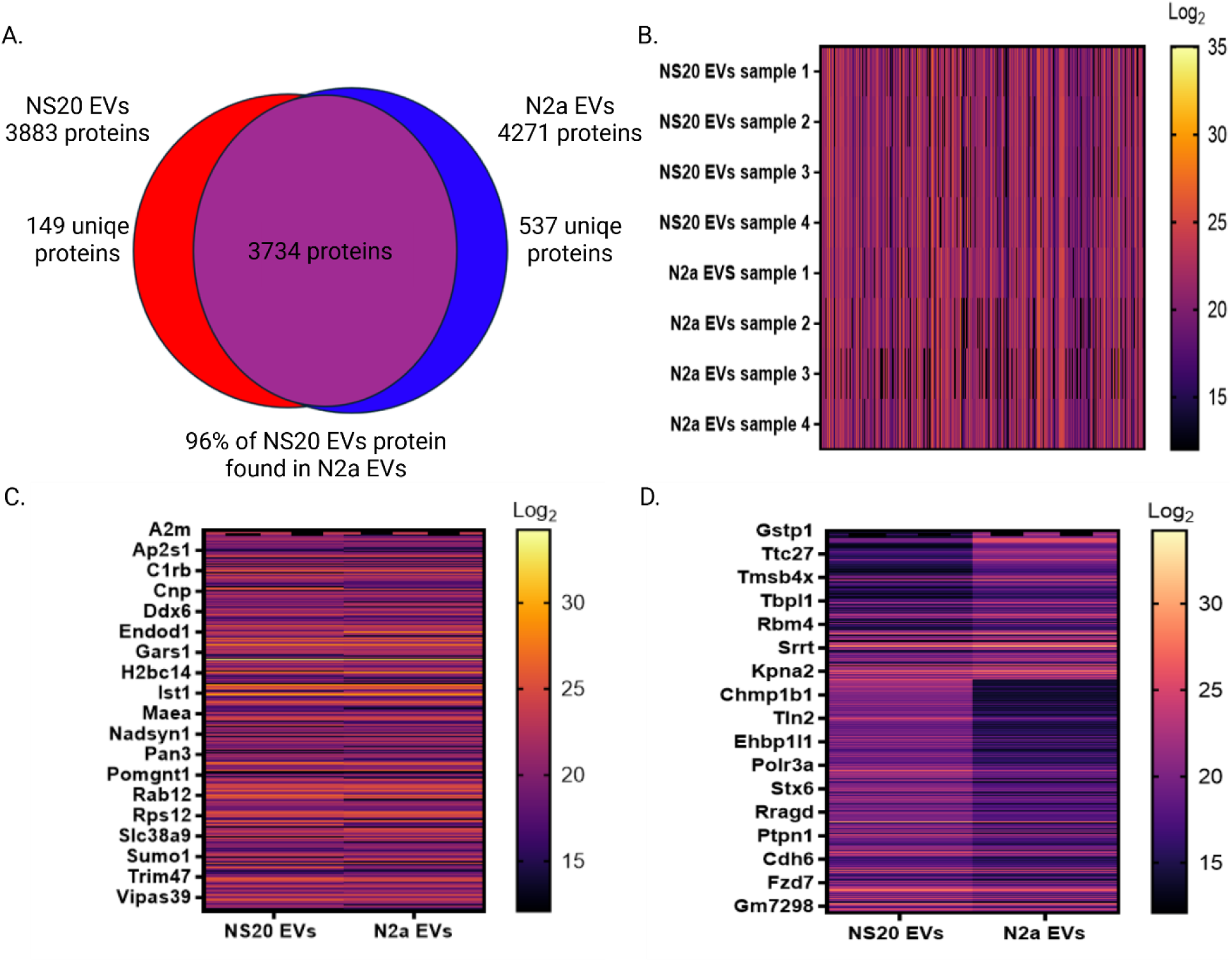
NB-EV proteomes are enriched in proteins involved in cell junction organization and adhesion. We found 3883 proteins in NS20, while 4271 proteins were found in N2a EVs. Additionally, 3734 proteins were found to be shared between the NB EVs, comprising 96% of all NS20 EV proteins (A). Heatmap of shared proteins between NS20 and N2a EVs (B). Mean log_2_ abundance of shared proteins (C). Among the 3734 proteins, 508 proteins were significantly upregulated (log_2_FC>1) in N2a EVs, while 794 were elevated in NS20 EVs (D).

### Proteomic analysis of NB-EVs cargo reveals tight-junction-associated proteins

Further analysis of the proteomics data using Cytoscape[34] identified neuroblast, neuronal, and neuroblastoma biomarkers in N2a and NS20 EVs. Neuroblast biomarkers such as DPYSL2, which plays a regulatory role in neurogenesis[35], and ST8SIA4, which produces polysialic acid (PSA)[36] for NCAM, were found in NB-EVs, which together may play a role in neuroblast migration and growth[37,38]. Additionally, we detected neuronal markers, such as MAP2, which plays a role in microtubule stabilization and regulates microtubule networks in axons and dendrites of neurons[39]. TUBB3, also found to be enriched in our proteomic data, plays a role in the neuronal cytoskeleton during neurogenesis[40]. VAMP2, a key protein in neuronal stability and synaptic vesicle fusion, has been shown to play a role in α-synuclein interactions in dementia[41]. While NCAM is a biomarker for neuroblast cells, it is also used as a biomarker for NB due to its overexpression at both the protein and RNA levels[42]. Vim, another cell adhesion protein that correlates with NB progression[43], was also detected in our NB EVs, as was Pes1, which is essential for NB cell survival and has been shown to be a promising prognostic biomarker[44]. In conclusion, 147 proteins (60 neuronal, 57 neuroblast/neurodevelopment, 30 NB) were enriched in N2a EVs (Figure 3A), and 122 proteins (57 neuronal, 51 neuroblast/neurodevelopment, and 14 NB) were enriched in NS20 EVs (Figure 3B). Among the shared neuronal proteins using a cutoff of log_2_FC>1, 32 were shared between both lines (4 elevated in NS20, 14 in N2a), 36 neuroblast/neurodevelopmental proteins were shared (13 elevated in NS20, 7 in N2a), and 11 NB (2 elevated in NS20, 4 in N2a) (Figure 3C). Proteomic data were examined for N2a EV proteins, cell junctions, cell junction assembly, and tight junction proteins, resulting in 1116 EV-associated proteins (53 were found thorough cytoscape, the rest are shared with NS20 EV associated proteins that were found in cytoscape), 271 cell junction assembly proteins, and 896 cell junction proteins being found, totaling 22.2% of the protein population found is associated with cell junctions and assembly (Figure 3D). In NS20 EVs, out of 3883 proteins, 1131 EV-associated proteins, 148 cell junction assembly proteins, and 856 cell junction proteins were identified, totaling 23% of the protein population associated with cell junctions (Figure 3E). Among the identified junction proteins, prominent proteins such as Occludin, a transmembrane TJ protein, are elevated in brain and CNS barrier endothelial cells[45]. Claudin-5, the most enriched tight junction protein, was also identified as highly enriched in NB EVs[46]. In addition, ZO-1 and ZO-2, essential TJ scaffold proteins that organize claudins and other junctional components into functional junctional strands[47], were also found in NB-EVs. On the same note, MMP-2 and MMP-9 were also identified in NB EVs. MMPs are essential in extracellular matrix (ECM) remodeling[48], and in CNS barriers[49,50] play a direct role in BBB leakage[51] via TJ disruption by degradation of Occludin and Claudin-5[52]. While N2a EVs encapsulated these TJ proteins and MMPs, we observed a significant enrichment of these proteins in NS20 EVs, in which Occludin, Claudin-5, MMP-2, and MMP-9 had elevated levels (133.422-fold, 93.574-fold, 3.312-fold, and 2.655-fold, respectively). N2a EVs, on the other hand, had more prevalent enrichment of ZO-1 (by 5.286-fold) and ZO-2 (by 1.223-fold) (Figure 3F).

**Figure 3.**
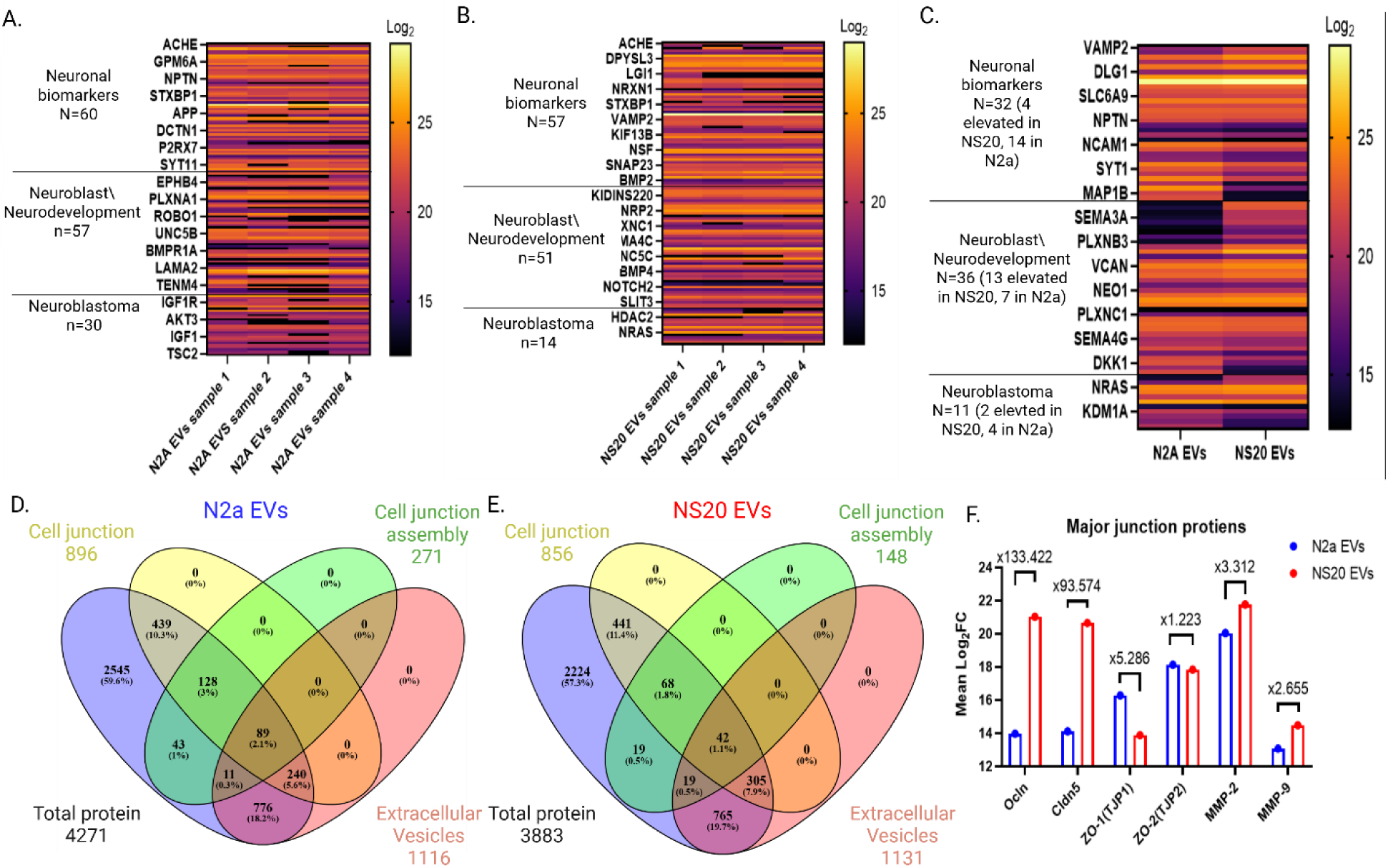
Proteomic analysis of NB EVs cargo reveals tight-junction-associated proteins. N2a EV proteomics (A). NS20 EV proteomics (B). Shared neuronal markers with a log2FC cutoff (C). N2a EVs show that, of 4271 proteins, 271 are associated with cell junction assembly, 896 with cell junctions, and 1116 are EV-associated proteins, totaling 22.2% of proteins associated with cell junction pathways (D). NS20 EVs show that out of 3883 proteins, 148 proteins are associated with cell junction assembly, 856 with cell junctions, and 1131 EV-associated proteins, totaling 23% of proteins that are associated with cell junction pathways (E). Occludin, Claudin-5, and MMPs 2 and 9 have been shown to be enriched in NS20 EVs, while ZO-1 and ZO-2 are more enriched in N2a EVs (F).

### NB-EVs carry functionally active MMPs

Proteomic identification of MMP-2 and MMP-9 within NB-derived EVs suggested that these vesicles may exert direct proteolytic effects that may be relevant to CNS barrier modulation. Since MMPs 2 and 9 are classified as gelatinases[53,54], we evaluated whether EV-associated MMPs are enzymatically active by gelatinase activity through a fluorescence-based DQ-gelatin assay. To minimize photobleaching associated with repeated fluorescence measurements[55,56], separate wells were assigned to different time-point sampling schemes. As shown in supplementary Figure 2A, EVs derived from both N2a and NS20 cells induced a significant increase in fluorescence over time, indicating substrate cleavage and active proteolysis. In contrast, negative control wells exhibited only minimal baseline signal, confirming that the observed fluorescence increase reflects EV-associated enzymatic activity. After normalization to background signal (supplementary Figure 2B), NB-EVs displayed a clear time-dependent increase in gelatinase activity, with a pronounced rise between 24 and 48 hours. To allow direct visualization of EV-derived activity, the positive control signal was omitted from subsequent plots due to its substantially higher intensity (supplementary Figure 2C). Quantitative analysis of fluorescence kinetics (Figure 4A) revealed that both N2a- and NS20-derived EVs exhibit robust proteolytic activity, with the most prominent increase occurring within the first 24 hours following exposure. Together, these results provide functional evidence that NB-EVs harbor enzymatically active gelatinases, supporting a mechanistic role for EV-associated MMPs in extracellular matrix remodeling and TJ destabilization. In addition, Zymography of NS20-EVs revealed bands at 72 kDa, which may represent the pro-active form of MMP-2[57] and above it, which may show the active form of MMP-9 at 82-85 kDa [58,59]. These results show that NB-EVs harbor forms of MMPs 2 and 9, which give them their proteolytic abilities (Figure 4.B).

**Figure 4.**
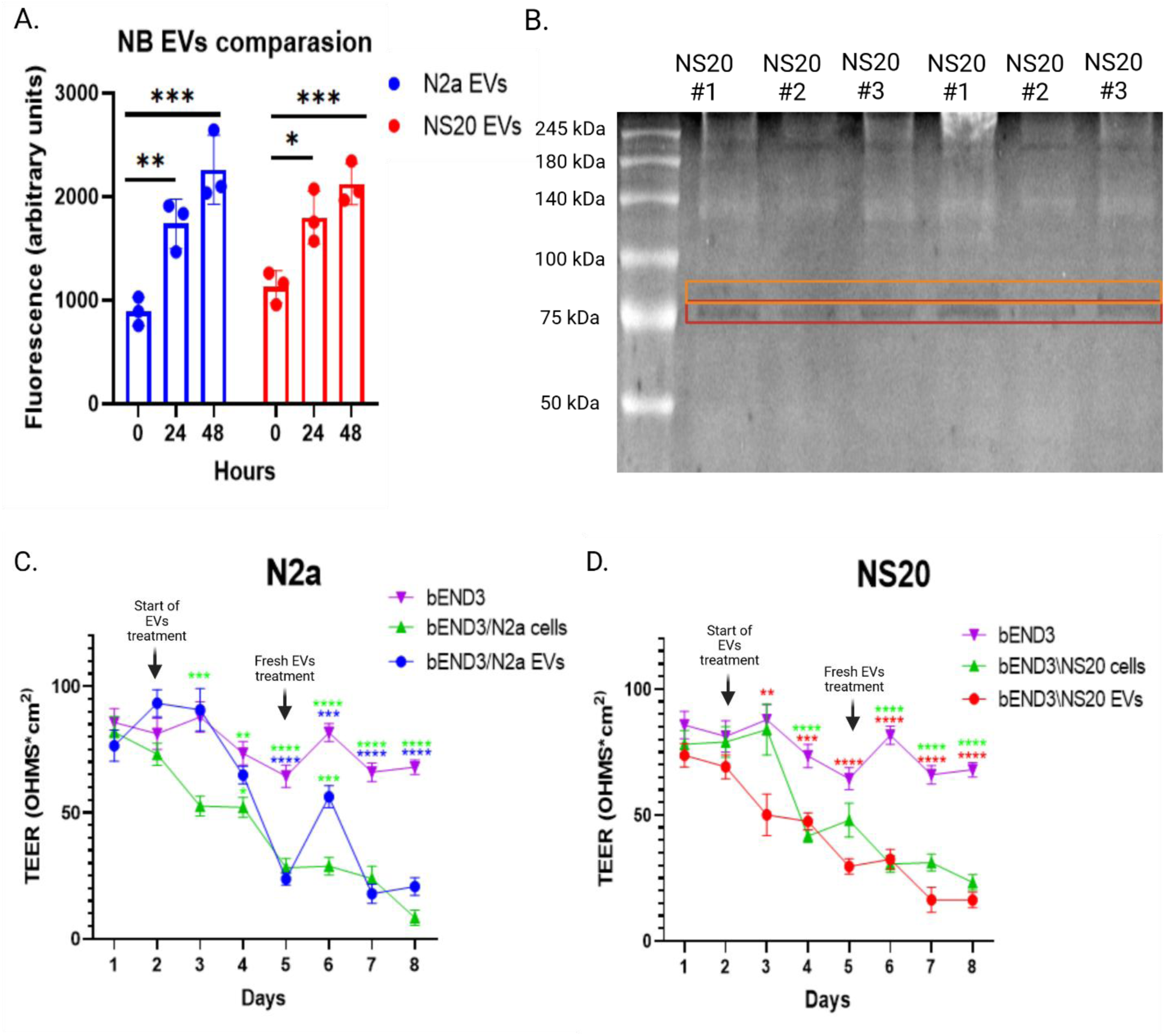
NB-EVs harbor functionally active MMPs. NB-EVs proteolytic activity was measured for 0,24,48 hours and measured (A). Zymography for NS20 EVs shows pro-active MMP-2 (red) and active MMP-9 (orange) (B). bEND3 monolayer TEER levels for 8 days after daily treatment with N2a EVs (C). bEND3 monolayer TEER levels for 8 days after daily treatment with NS20 EVs (D). We analyzed MMP activity using an ordinary two-way ANOVA with cell line and time as fixed factors. We did not apply a repeated-measures structure because measurements at each time point were obtained from independent wells. We used a Tukey post hoc multiple-comparisons test to evaluate pairwise differences across time points within each cell line and to compare cell lines at matching time points. Data are presented as mean ± SEM, and multiplicity-adjusted P-values are reported for all comparisons. We defined statistical significance as P < 0.05. For TEER experiments in which the same wells were measured over eight consecutive days, we performed a two-way repeated-measures ANOVA, with Time as the within-subject factor and Group as the between-subject factor. Matched measurements were stacked into sub columns to account for repeated observations per well. Because sphericity could not be assumed, we applied the Geisser–Greenhouse correction. We performed post-hoc comparisons using Tukey’s multiple-comparisons test, evaluating group differences at each time point (row-wise comparisons, one family per row). Statistical significance was indicated as follows: *p < 0.05, **p < 0.01, ***p < 0.001, ****p < 0.0001. All TEER experiments were performed with 3 biological replicates, 6 reads per biological replicate per group to ensure robustness and reproducibility.

To test whether NB-derived EVs may functionally alter CNS endothelial barrier integrity, we used a commonly used in vitro model[60–62]. Murine brain endothelial cells (bEND3) were seeded as a monolayer on a transwell insert. This system permits functional assessment of TJ formation by monitoring trans-endothelial electrical resistance (TEER), which correlates with TJ integrity in endothelial cells[63–66]. Following exposure to EVs derived from N2a or NS20 cells. As shown in Figure 4C, repeated exposure to N2a-derived EVs resulted in a significant decline in TEER overtime, indicating a gradual loss of endothelial tight-junction integrity. While an initial exposure to freshly isolated EVs transiently increased TEER, continued EV treatment led to a sustained, pronounced disruption of the barrier. Co-culture with N2a tumor cells induced a similar, albeit more pronounced, decline in TEER, suggesting that EVs contribute substantially to tumor-mediated barrier impairment. NS20-derived EVs elicited a more rapid and sustained reduction in TEER compared with N2a EVs (Figure 4D). Barrier disruption was evident within 24 hours of initial exposure and progressed steadily throughout the experiment, with no evidence of recovery following exposure to freshly isolated EVs. Co-culture with NS20 cells similarly destabilized endothelial integrity, supporting the notion that mature neuroblast NB cells release EVs with heightened barrier-disruptive capacity. Quantitative comparison revealed that both N2a and NS20-derived EVs induced TEER reductions approaching those observed with direct tumor cell co-culture, underscoring the potency of EVs in modulating endothelial function.

### NB-EVs disrupt CNS endothelial barrier integrity via MMP-dependent mechanisms

Given our findings in Figures 3 and 4, which suggest the presence of enzymatically active MMP-2 and MMP-9 in NB-EVs, we next examined whether EV-induced barrier disruption is mediated by metalloproteinase activity. To that end, we used pharmacological MMP inhibition with the broad-spectrum zinc chelator 1,10-phenanthroline. Notably, this inhibitor prevented the EV-induced TEER decline for both N2a (Figure 5A) and NS20-derived EVs (Figure 5B). In the presence of MMP inhibition, TEER values remained substantially closer to untreated controls, indicating a significant preservation of endothelial barrier integrity. Importantly, 1,10-phenanthroline alone exerted only a modest effect on TEER, suggesting that its protective effect primarily reflects inhibition of EV-associated proteolytic activity rather than nonspecific toxicity. To determine whether the effects of NB-EVs on bEND3 endothelial cells represent a tumor-associated phenotype rather than a general response to EV exposure, we compared NB-EVs with EVs derived from non-tumor control cells, including astrocytes and fibroblasts. Astrocytes comprise a major non-neuronal cell population in the neurovascular unit. Traditionally, in the CNS, astrocytes are known to be involved in homeostasis, management of extracellular ion balance and pH, regulation of neurotransmission, and control of cerebral blood flow[67]. Although fibroblasts are not a direct component of the neurovascular unit, they have been shown to support endothelial and epithelial TEER levels when co-cultured[68]. In addition, fibroblasts have recently been studied for their role in the neurovascular unit and CNS [69,70]. Thus, we again established bEND3 monolayers for TEER measurements, but instead of the NB-EVs, we used C8D1A (astrocyte) and NIH-3T3 (fibroblast) EVs for 3 days with the same daily dose as the NB-EVs and 1,10-phenanthroline. The TEER measurements show that C8D1A (Figure 5C) and NIH-3T3 (Figure 5D) increased TEER levels compared with NB-EVs, which showed the opposite effect. 1,10-phenanthroline didn’t show a significant increase compared with the non-tumor EVs. Together, these data demonstrate that NB-EVs are sufficient to disrupt CNS endothelial barrier function and that this effect is at least partially dependent on EV-associated MMP activity. The distinct kinetics observed between early- and late-differentiation NB-EVs further support the existence of stage-dependent EV programs that impair CNS barrier integrity by distinct, yet convergent mechanisms. In addition, the effects that NB-EVs have shown did not repeat themselves; on the contrary, they showed that EVs of supporting cells increase the TEER levels, which have been shown in previous work on astrocyte EVs[71].

**Figure 5.**
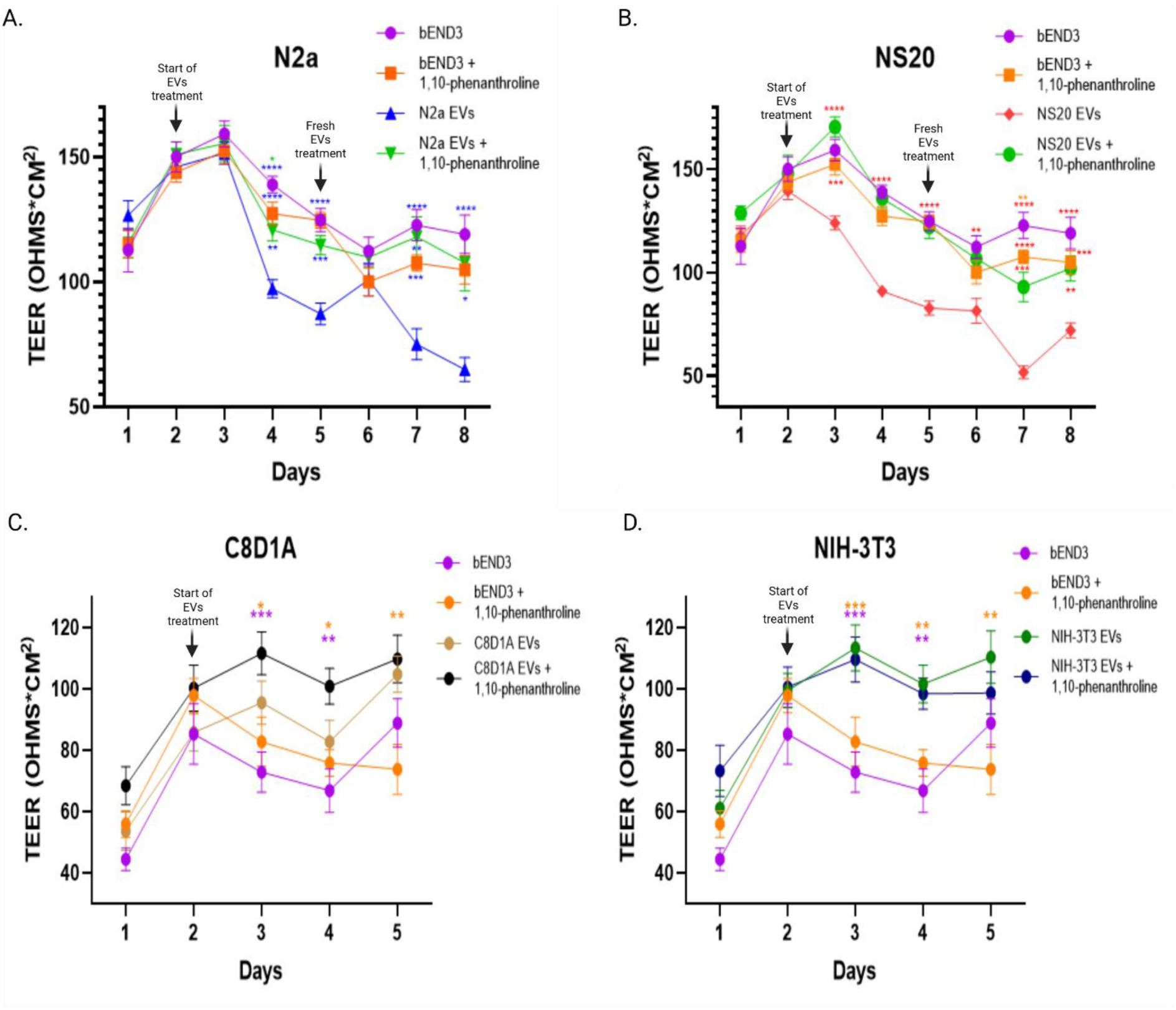
NB-EVs disrupt CNS endothelial barrier integrity via MMP-dependent mechanisms. A repeated setup was established; in addition, 0.2mM 1,10-phenanthroline was added as an MMP inhibitor to N2a EVs (A) and NS20 EVs (B). We repeated the setup for 5 days using EVs derived from the astrocytic cell line C8D1A (C), and EVs derived from the fibroblast cell line NIH-3T3 (D). Statistical analyses were conducted and presented as mean ± Standard Error of the Mean (SEM) unless otherwise specified. For TEER experiments in which the same wells were measured over eight consecutive days, we performed a two-way repeated-measures ANOVA, with Time as the within-subject factor and Group as the between-subject factor. Matched measurements were stacked into subcolumns to account for repeated observations per well. Because sphericity could not be assumed, we applied the Geisser–Greenhouse correction. We performed post-hoc comparisons using Tukey’s multiple-comparisons test, evaluating group differences at each time point (row-wise comparisons, one family per row). Statistical significance was indicated as follows: *p < 0.05, **p < 0.01, ***p < 0.001, ****p < 0.0001. All TEER experiments were performed with 3 biological replicates, 6 reads per biological replicate per group to ensure robustness and reproducibility.

## Discussion

NB remains a clinically challenging pediatric malignancy characterized by pronounced heterogeneity, frequent relapses in high-risk disease, and significant neurological morbidity[72]. While tumor cell– intrinsic drivers of NB progression have been extensively studied, far less is known about how NB cells communicate with and remodel distant microenvironments, particularly within the CNS. In this study, we identify EVs released from NB cells as potential mediators of CNS endothelial barrier dysfunction and provide mechanistic evidence linking EV cargo composition to TJ destabilization and proteolytic activity. Our functional data first demonstrates that NB tumor cells themselves exert a pronounced disruptive effect on CNS endothelial barrier integrity, as evidenced by the marked TEER decline observed during insert co-culture with both early (N2a) and late (NS20) neuroblastoma cells. This decline in TEER indicates that indirect extracellular communication contributes to that phenotype, which correlates with our data demonstrating that EVs released from both early (N2a) and late (NS20) NB cells significantly impair endothelial barrier integrity, as functionally assessed by TEER in a well-established murine CNS endothelial model[73]. Notably, although both EV populations induced substantial TEER reduction, the kinetics and reversibility of barrier disruption differed between early- and late-differentiated NB-EVs. N2a-derived EVs induced a delayed yet pronounced disruption that exhibited transient recovery upon exposure to fresh EVs, whereas NS20 EVs elicited a more immediate and sustained decrease in TEER without evidence of recovery. These differences likely reflect stage–specific EV programs, suggesting that NB neuro-progression is accompanied not only by changes in EV quantity or size but also by qualitative shifts in functional cargo.

Proteomic profiling of NB-EVs revealed a striking enrichment of proteins associated with cell junction organization, adhesion, and structural integrity. Approximately one-quarter of the total EV protein cargo in both N2a and NS20 EVs was linked to junctional pathways, including canonical TJ proteins such as Occludin and Claudin-5. Claudin-5 is a key determinant of endothelial barrier selectivity in CNS vasculature, and its dysregulation has been directly implicated in BBB leakage in both pathological and experimental settings[74]. The presence of these junctional components within EVs suggests a dual mechanism by which NB-EVs may disrupt barrier integrity: direct delivery of junction-associated proteins that can interfere with endothelial junction assembly, and indirect enzymatic degradation of endogenous junctional complexes.

Crucially, our proteomic analysis also identified matrix metalloproteinases MMP-2 and MMP-9 within NB-EVs, which were also found via zymography. These gelatinases are well-established mediators of BBB breakdown and cleave TJ proteins, including Occludin and Claudin-5[75,76], thereby increasing paracellular permeability. Functional assays confirmed that NB-EVs possess robust gelatinase activity, peaking within 24 hours of exposure. The rescue of TEER levels upon MMP inhibition indicates that proteolysis represents a major, but not exclusive, mechanism underlying EV-mediated barrier dysfunction. Since inhibition provides TEER rescue, the data argue that the effect is predominantly MMP-driven (likely MMP-2/9; possibly also non-gelatinase MMP-11/12). Other proteases present in the EV proteome (ADAM/ADAMTS, serine proteases, cathepsins) are less likely to be major contributors to the TEER phenotype in this assay, at least under our conditions. Thus, NB-EVs likely engage multiple complementary pathways to destabilize CNS barriers, collectively lowering the threshold for tumor dissemination and neuroinflammation. These results have also been supported via C8D1A and NIH-3T3 EVs TEER measurements, which showed an increase in TEER levels while the NB-EVs showed a rapid decline, thus showing that the tumor-oriented EVs are playing an invasive role in comparison to the non-tumor-supporting cells EVs.

From a disease perspective, these findings have important implications for NB dissemination and CNS involvement. BBB disruption by tumor-derived EVs may facilitate tumor cell extravasation into the CNS, increase neural tissue exposure to circulating tumor-derived factors, and contribute to neurological dysfunction and seizures even in the absence of overt tumor metastases[77,78]. The presence of neuronal and neuroblast-specific markers within EVs further suggests that these vesicles may participate in pre-metastatic niche formation or modulate neural circuits to favor tumor survival and progression.

This study has several limitations. Primarily, our work relies on murine NB cell lines and an in-vitro endothelial model, which cannot fully recapitulate the cellular complexity and hemodynamic forces present in vivo. However, an important advantage of such reductionist in vitro systems is the ability to selectively couple and uncouple individual components, namely tumor-derived EVs, endothelial cells, and neuroblastoma cells, thereby enabling precise mechanistic dissection of their direct and combinatorial contributions to BBB dysfunction. This modular control represents a recognized strength of advanced in vitro BBB platforms, including organ-on-chip systems that bridge in vitro and in vivo biology[79]. Future studies employing selective inhibition, genetic perturbation, or in vivo models will be required to delineate the relative contributions of specific EV-associated proteases and to validate the relevance of these mechanisms in the intact CNS. Additionally, extending these findings to human NB-derived EVs and patient samples will be critical for translational validation.

Nevertheless, proteomic analyses of circulating extracellular vesicles from NB patients, including both plasma exosomes[80] and serum small extracellular vesicles (sEVs)[81], have independently identified EV signatures enriched for extracellular matrix remodeling, adhesion, cytoskeletal organization, and tumor progression pathways, with distinct profiles distinguishing high-risk from low-risk disease. These clinical datasets demonstrate that NB tumors release biologically meaningful EV populations into the circulation; however, they do not address the functional consequences of this cargo on target tissues. Our cell-derived EV data provides the missing mechanistic link by showing that NB-EVs are highly enriched in TJ components and proteolytically active gelatinases, and that they are sufficient to induce CNS endothelial barrier breakdown. The strong convergence between the ECM-, adhesion-, and invasion-associated pathways observed in patient-derived sEVs and the barrier-disruptive EV program identified in our N2a and NS20 models supports a unified model in which circulating NB-EVs detected in patients represent the in vivo manifestation of the same EV machinery that actively remodels tissue barriers. Together, these findings indicate that patient sEV proteomic signatures are not merely biomarkers of disease state but reflect a biologically active EV-mediated mechanism that contributes directly to CNS vulnerability, metastatic dissemination, and neurological morbidity in NB.

## Conclusion

In summary, our work identifies NB-derived EVs as central regulators of CNS barrier integrity and uncovers a mechanistic link between EV-associated TJ proteins, proteolytic activity, and endothelial dysfunction. These findings expand current understanding of NB–CNS interactions and highlight EVs as potential therapeutic targets to limit barrier disruption and metastatic progression. Targeting EV biogenesis, cargo loading, or EV-associated proteases may therefore represent novel therapeutic strategies to preserve CNS barrier function, limit metastatic dissemination, and mitigate neurological complications in NB patients. More broadly, this work underscores the importance of tumor-derived EVs as dynamic modulators of tissue barriers and highlights their potential as both biomarkers and therapeutic targets in pediatric cancer.

## List of abbreviations

ADAM: A disintegrin and metalloproteinase
ADAMTS: A disintegrin and metalloproteinase with thrombospondin motifs
ANOVA: Analysis of variance
BBB: Blood–brain barrier
BTB: Brain–tumor barrier
CNS: Central nervous system
DMEM: Dulbecco’s Modified Eagle Medium
ECM: Extracellular matrix
EV: Extracellular vesicle
FBS: Fetal bovine serum
MMP: Matrix metalloproteinase
NB: Neuroblastoma
NK: Natural killer
NTA: Nanoparticle tracking analysis
OMAS: Opsoclonus-myoclonus-ataxia syndrome
PBS: Phosphate-buffered saline
SEC: Size-exclusion chromatography
SEM: Standard error of the mean
SNS: Sympathetic nervous system
TEER: Trans-endothelial electrical resistance
TEM: Transmission electron microscopy
TJ: Tight junction
TME: Tumor microenvironment
ZO: Zonula occludens

## Declarations

### Ethics approval and consent to participate

Not applicable.

### Consent for publication

Not applicable.

### Availability of data and materials

The datasets generated and/or analyzed during the current study are available from the corresponding author on reasonable request.

### Authors’ contributions

T.M.: Conceptualization, methodology, investigation, data curation, formal analysis, visualization, and writing – original draft. I.L.: Methodology, investigation, project administration and validation. M.Z.: Methodology, investigation, and validation. V.F.: Methodology and validation. S.B.D.: Methodology, investigation, and validation. E.B.-Y.: Methodology, review and editing. G.V.: Conceptualization, methodology, resources. T.C.: Conceptualization, supervision, project administration, funding acquisition, and writing – review and editing. All authors read and approved the final manuscript.

## Acknowledgements

N/A

