## Supplemental for "Neuroblastoma-derived Extracellular Vesicles Disrupt the Integrity of Central Nervous System Barriers via Tight Junction Modulation"

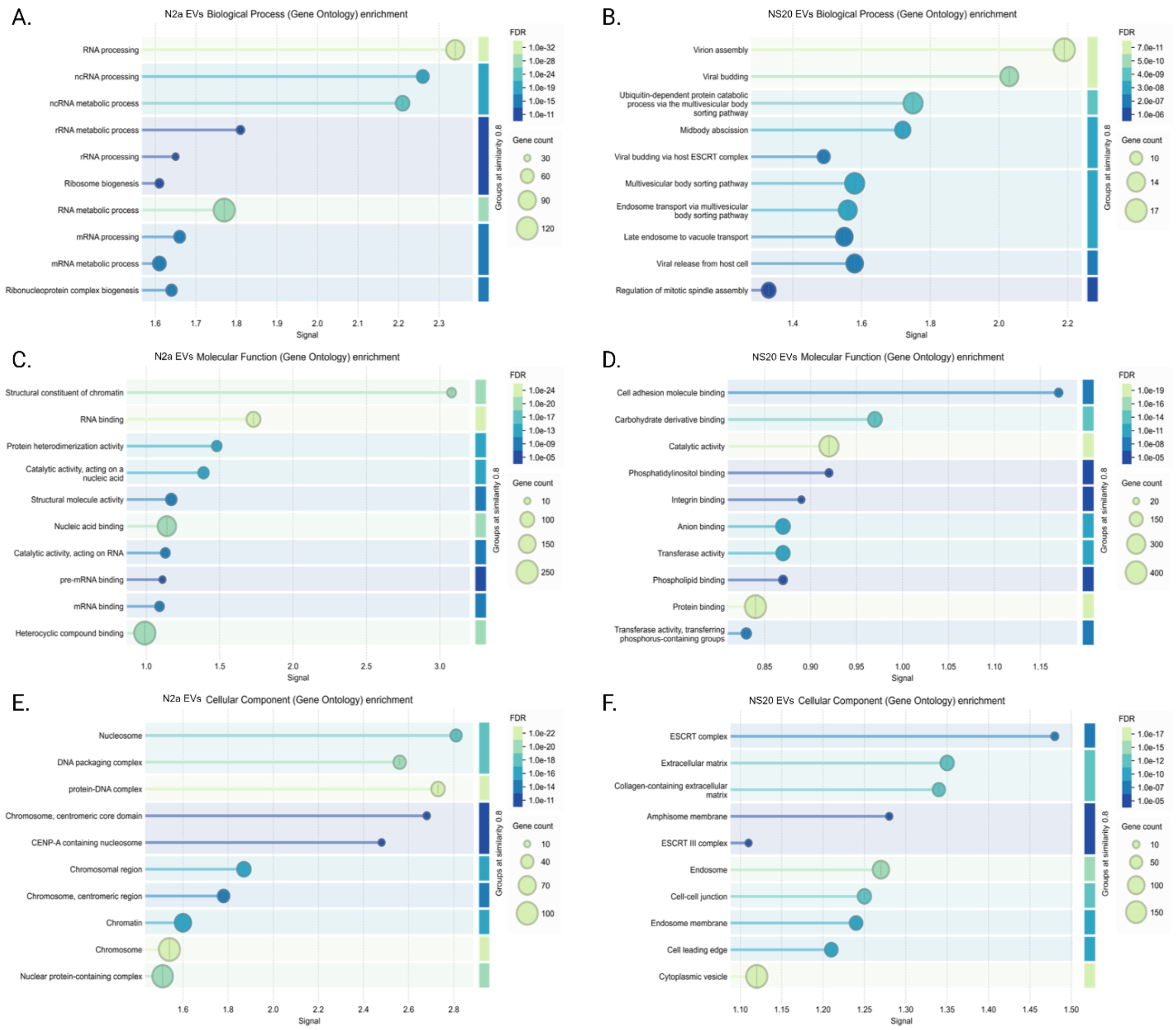

**Supplementary Figure 1 – STRING analysis for NB-EVs proteomics** - STRING analysis of significantly enriched proteins was performed using a high-confidence interaction score of 0.7. Functional enrichment analyses show biological process terms for N2a and NS20 EVs (A and B), molecular function terms (C and D), and cellular component terms (E and F), respectively.

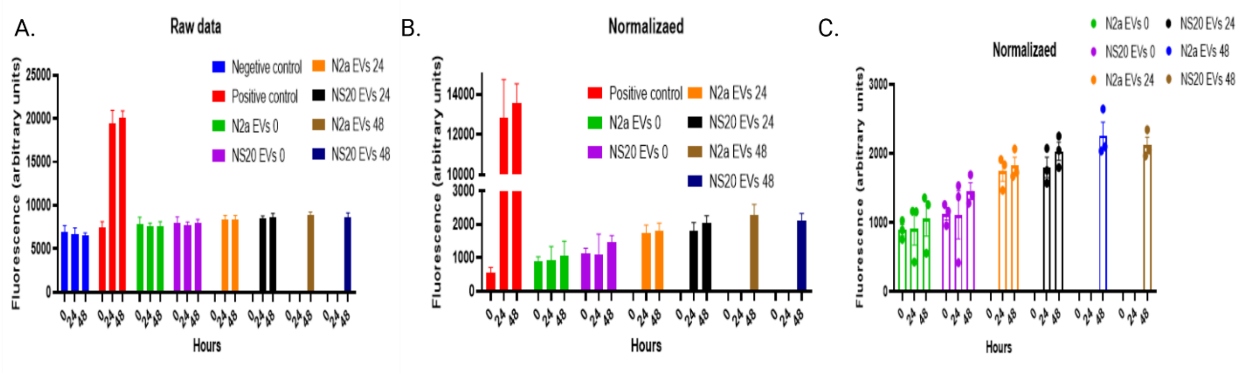


**Supplementary Figure 2 – proteolytic activity of NB-EVs setup –** Raw fluorescence measurements of the proteolytic activity of NB-EVs at 0, 24, and 48 h (A). Data normalized by subtraction of the negative-control background (B). Normalized NB-EV data shown without the positive control to improve visualization of EV-associated proteolytic activity (C).

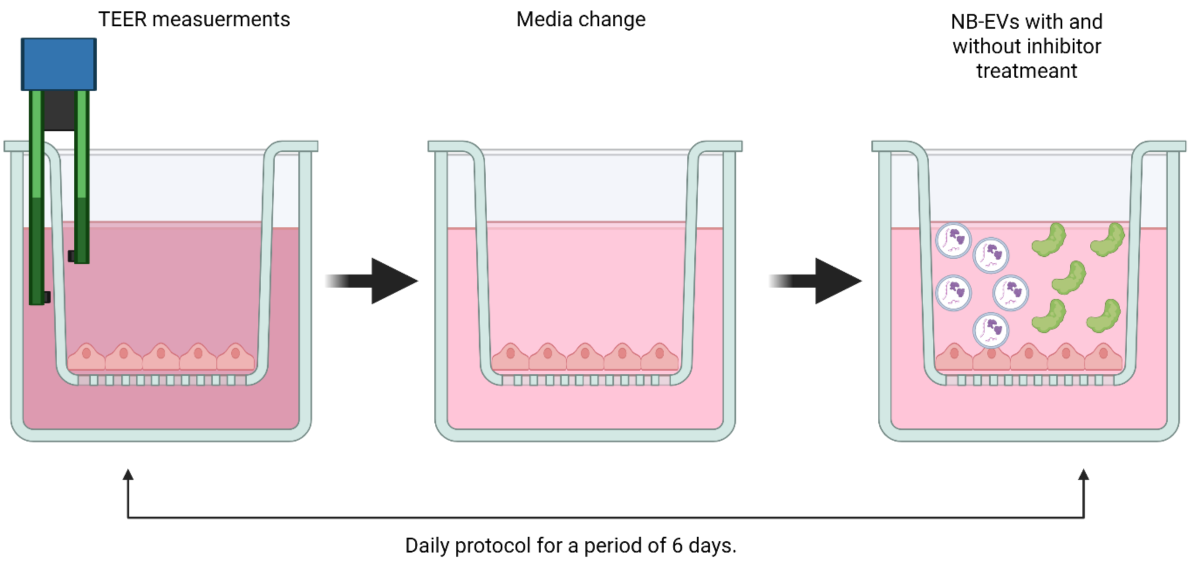


**Supplementary figure 3 – bEND3 TEER setup** – Following coating of 12-well Transwell inserts with collagen IV and fibronectin, bEND3 cells were seeded and cultured for 2 days before TEER measurements were initiated. Following each TEER measurement, the medium was replaced, and cells were treated with NB-EVs with or without an MMP inhibitor. This procedure was repeated daily for 6 days unless otherwise stated.
